# ggPlantmap: an R package for creation of informative and quantitative ggplot maps derived from plant images

**DOI:** 10.1101/2023.11.30.569429

**Authors:** Leonardo Jo, Kaisa Kajala

## Abstract

As plant research generates an ever-growing volume of spatial quantitative data, the need for decentralized and user-friendly visualization tools to explore large and complex datasets tools becomes crucial. Existing resources, such as the Plant eFP (electronic Fluorescent Pictograph) browsers, have played a pivotal role on the communication of gene expression data across many plant species. However, although widely used by the plant research community, the Plant eFP browser lacks open and user-friendly tools for the creation of customized expression maps independently. Plant biologists with less coding experience can often encounter challenges when attempting to explore ways to communicate their own spatial quantitative data. We present ‘ggPlantmap’ an open-source R package designed to address this challenge by providing an easy and user-friendly method for the creation of ggplot representative maps from plant images. ggPlantmap is built in R, one of the most used languages in biology to empower plant scientists to create and customize eFP-like browsers tailored to their experimental data. Here, we provide an overview of the package and tutorials that are accessible even to users with minimal R programming experience. We hope that ggPlantmap can assist the plant science community, fostering innovation and improving our understanding of plant development and function.

**Highlight:** ggPlantmap, a new addition to the plant data visualization toolbox, allows users to create graphical maps from plant images for the representation of spatial quantitative data in R.

## Introduction

Over the last decade, the increased accessibility of sequencing technologies has allowed for a diverse array of plant research groups to generate sequencing data to address a variety of plant biology questions. More recently, development of single-cell and spatial sequencing techniques added a spatial dimension to this vast and growing mountain of data (Libault *et al*., 2017; Cuperus, 2022). These approaches enable the characterization of cell-specific events, allowing us to gain novel insights into the spatial organization of minute biological processes occurring in complex plant tissues with unprecedented resolution (Lee *et al*., 2023; Nobori *et al*., 2023). In addition to transcriptome analysis, plant scientists have successfully profiled proteins and metabolites within individual cell types, enabling a comprehensive understanding of the signaling pathways of specific cell types. We are entering a new and exciting paradigm of plant systems biology where we can now dissect individual cellular responses to a range of biotic and abiotic stimuli within highly heterogeneous tissues (Clark et al., 2022; Pandian et al., 2023). As these types of data are continuously being generated within the plant research community, there is a rising demand for the development of specialized data visualization tools that can effectively summarize, explore, and communicate cell or tissue specific quantitative data.

The ability to visualize the spatial distributions of gene expression, hormone level, signaling response, or other quantitative data on representative maps of tissue geometries can be an effective way to communicate the intricacies of highly complex datasets (Figure 1A). Spatial quantitative data, when represented on top of graphical maps (Figure 1A), offer the observers an opportunity to recognize patterns in their data and to formulate hypotheses, enhancing the interpretability and insights derived from the data visualization. For instance, the Plant eFP (electronic fluorescent pictograph) browsers (Figure 1B) have been an extremely valuable resource for researchers seeking to visualize gene expression data in the context of plant tissues across many different plant species (Winter *et al*., 2007; Waese *et al*., 2017; Waese-Perlman *et al*., 2021). The ability to display gene expression profiles in plant eFP browsers (Figure 1B), has proven to be a highly effective approach for visualization, communication, and education within the plant sciences. Not surprisingly, the Plant eFP browser has been one of the most cited plant resources to date, and the platform has been integrated in several comprehensive plant databases (Waese-Perlman *et al*., 2021). However, the development of plant eFP browsers for data generated from additional species, organs and tissues depends on the Bio-Analytic Resource for Plant Biology (BAR) team (https://bar.utoronto.ca/welcome.htm). Their efforts play a pivotal role in creating valuable resources that facilitate the exploration and understanding of plant transcriptome data. Over the years, the BAR group has been instrumental in supporting many plant research groups in creating unique plant eFP browsers for many different plant species. As more research groups generate and explore new transcriptome data, a single group of researchers generating visual maps for all this data will inevitable become unsustainable. It is therefore important to empower and equip plant researchers with tools to visualize, explore and navigate through their data independently. Allowing researchers to generate and customize their own eFP-like browsers can foster the creativity and ensure a more sustained progress of plant research. This decentralized approach, however, is challenged due to technical difficulties in generating quantitative displaying maps in an efficient, user-friendly manner. Experimental plant scientists, although more and more trained in scripting for basic data analysis, bioinformatics or visualization purposes are typically not apt programmers and therefore likely to encounter substantial challenges when attempting to incorporate and further explore their own spatial quantitative data. Alternatively, graphic software (i.e., Adobe Illustrator, Microsoft PowerPoint) can be used to manually map quantitative data on top of graphical images of tissue geometry. However, this approach is non-automatic and hence neither scalable nor suitable for exploring large datasets within coding pipelines.

**Figure 1.**
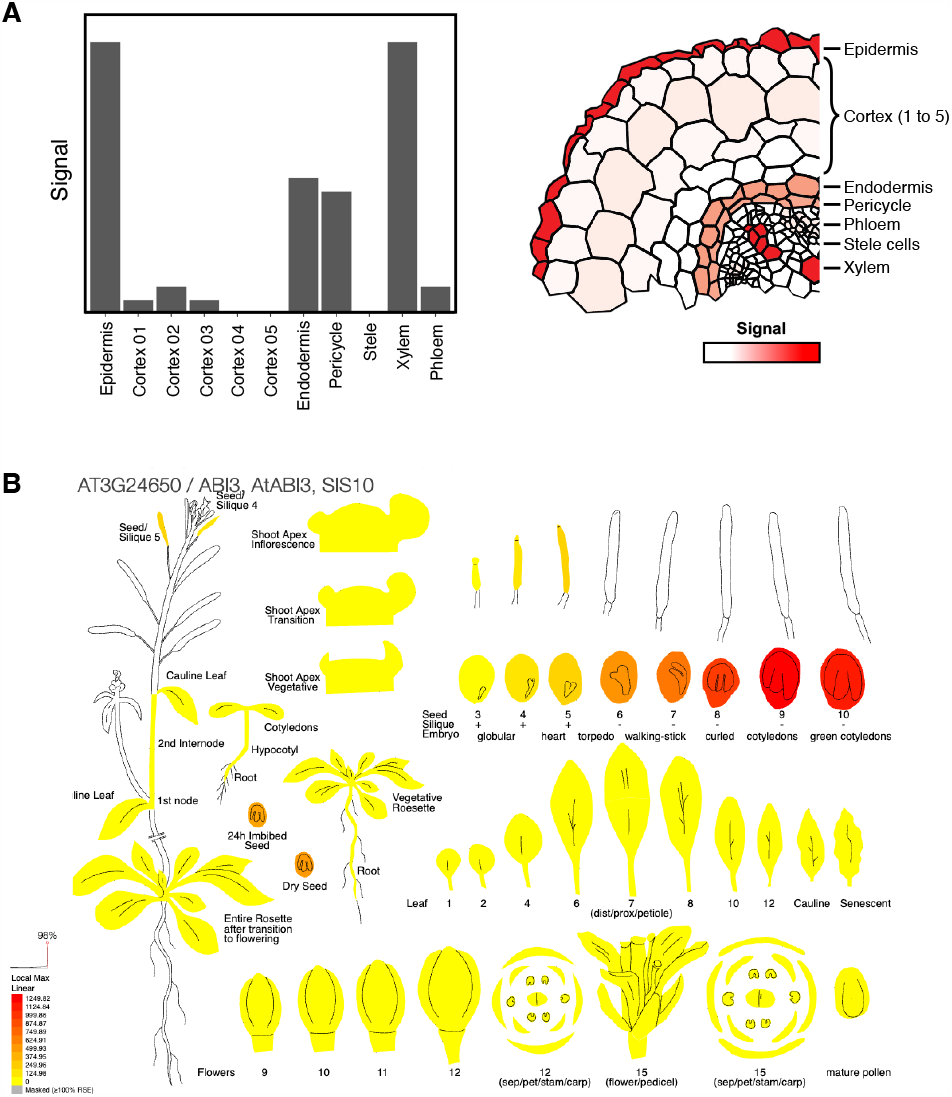
Different ways of displaying quantitative data in distinct plant tissues. **(A)** Two different ways to graphically represent a hypothetical quantitative signal across different cell types of the *Medicago sativa* root. On the left panel, values are plotted as conventional bar plots. On the right panel, values are represented as a heatmap on top of a representative cross-section image of the root. Note that in the right image, readers can rely on the spatial mapping of the signal to identify peculiarities in a quantitative pattern. **(B)** Plant eFP browser showing the expression of ABI3 (*AT3G24650*) in distinct Arabidopsis tissues. Image extracted from the Plant eFP browser (https://bar.utoronto.ca/eplant/).

To fill the existing resource gap, we introduce ‘ggPlantmap’, an open-source R package with the goal of providing a simple and easy way to create eFP-like visualizations using the R language (Figure 2). Since R is one of the most versatile programming languages and is widely used among biologists, we believe that this package is both accessible and useful for a wide range of users. We hope to inspire and empower potential users to use ggPlantmap to create visualizations of any quantitative data on any map (diagram). Here, we present an overview of the package functions and step-by-step guide to help users create new map diagrams from plant images and to integrate external quantitative data to create representative heatmaps (Figure 2). With the guide provided here as well as the documented guides found in the package website, we hope to foster the independent usage of the package by the plant research community.

**Figure 2.**
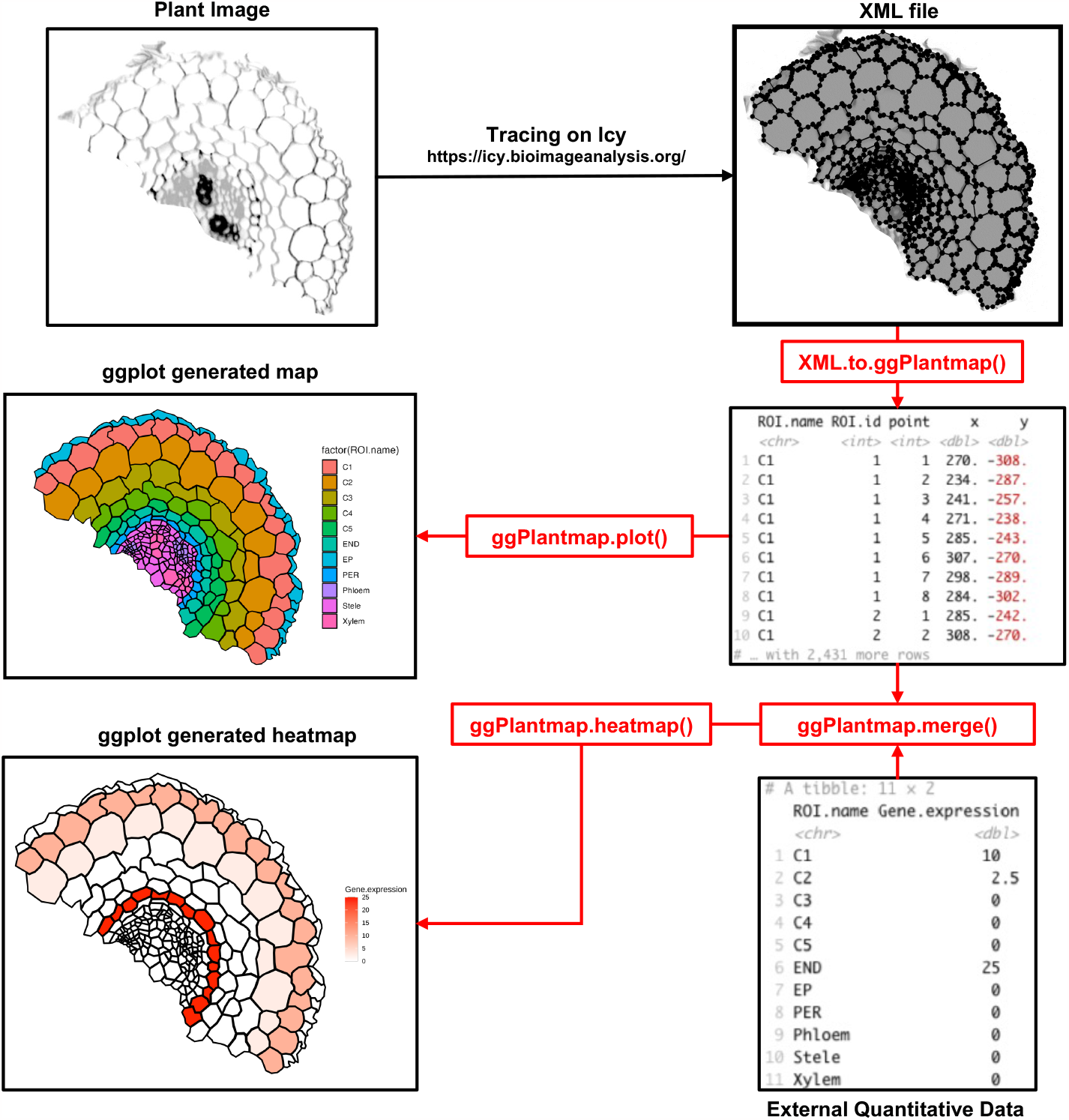
Overview of the ggPlantmap functions. Plant images are traced using the Icy software (https://icy.bioimageanalysis.org). These traces are stored as XML files and converted into map tables using the XML.to.ggPlantmap() function. These tables can be used to produce a ggplot using the ggPlantmap.plot() function or to be combined with external quantitative data to produce a ggplot heatmap using the ggPlantmap.merge() and ggPlantmap.heatmap() functions.

### Overview of the ggPlantmap package

The overview of the package functions is summarized in Figure 2. Plant images are manually traced using Icy, the open-source software for image analysis (https://icy.bioimageanalysis.org/) (De Chaumont *et al*., 2012). The traced polygons are stored as Extensible Markup Language (XML) files that can later be converted into map table objects using the XML.to.ggPlantmap() function. The structure of a ggPlantmap table emulates existing geographic map table formats in R, with a set of x and y coordinates of polygon points in a cartesian space (Figure 3). By relying on a known and simple table structure, the package ensures ease of use for users familiar with the R language. Additionally, we believe that with this simple and well-known table format, users can share their custom maps in a consistent and standardized manner, fostering efficient communication and collaboration within the plant scientific community.

**Figure 3.**
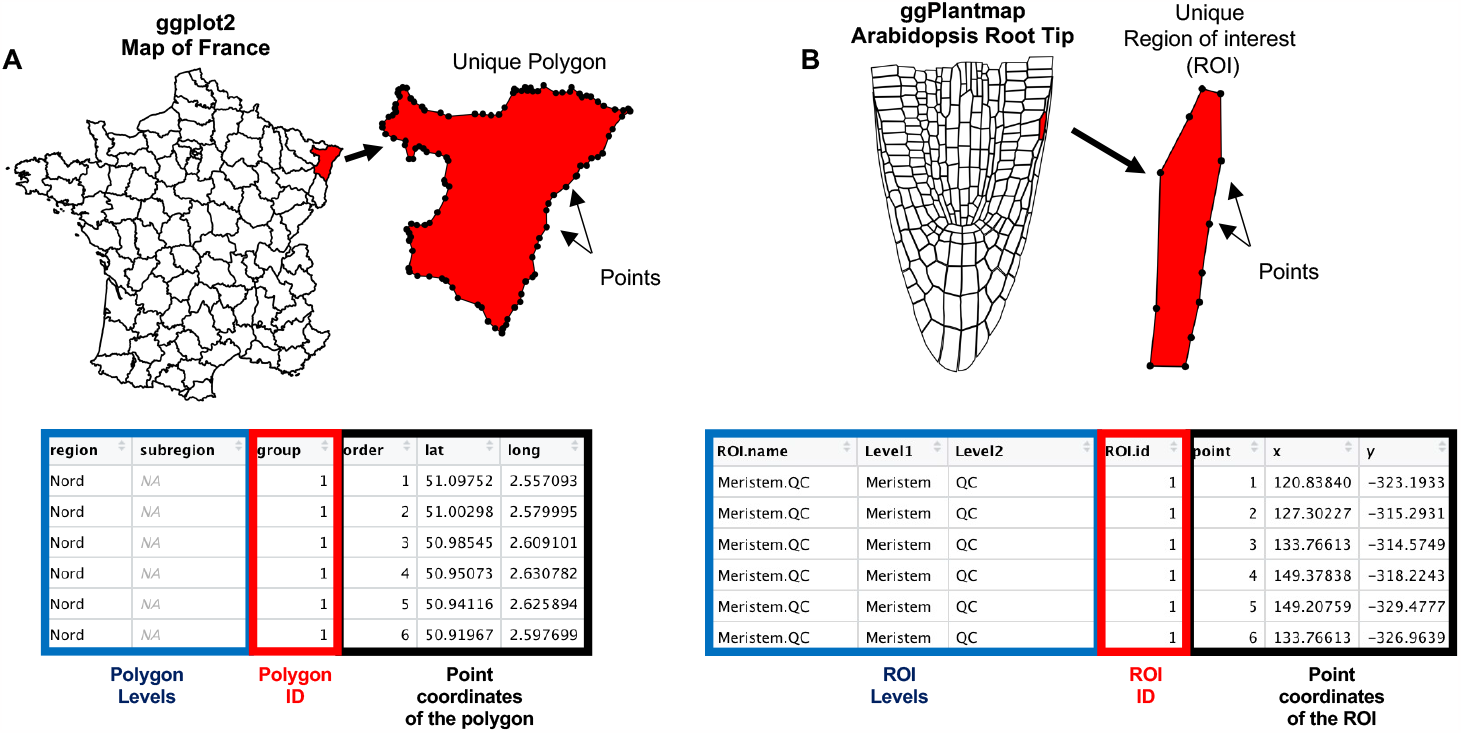
ggPlantmap map format is similar to ggplot2 geographical map table structure. In ggplot2, maps (A) are defined as combinations of polygons. These polygons are stored in tables with latitudinal (lat) and longitudinal (long) coordinates of unique points of each specific polygon. Similarly, in ggPlantmap **(B)**, distinct region of interest (ROI) is delimited by unique points with x and y coordinates in a Cartesian space.

The package includes a set of map tables created from previously published plant images that can be directly inserted into a R coding workflow (Table 1, Figure 4). These pre-loaded map tables are automatically loaded into the R environment during installation and loading of the ggPlantmap (library(ggPlantmap)) and their detailed information can be retrieved in the pre-loaded ggPm.summary table object.

**Table 1.**
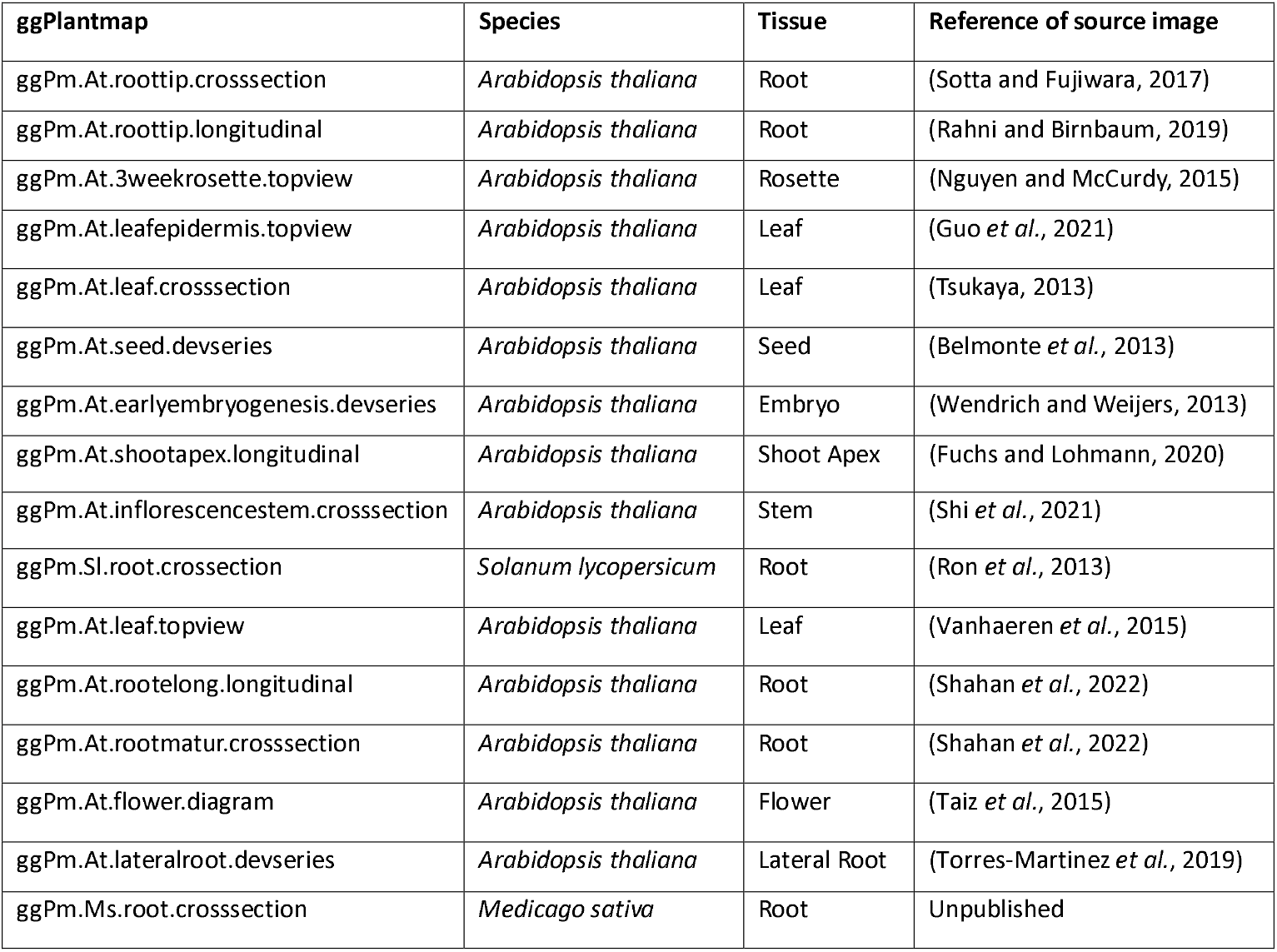
List of pre-loaded map tables in the ggPlantmap package.

| ggPlantmap | Species | Tissue | Reference of source image |
| --- | --- | --- | --- |
| ggPm.At.roottip.crossection | *Arabidopsis thaliana* | Root | (Sotta and Fujiwara, 2017) |
| ggPm.At.roottip.longitudinal | *Arabidopsis thaliana* | Root | (Rahni and Birnbaum, 2019) |
| ggPm.At.3weekrosette.topview | *Arabidopsis thaliana* | Rosette | (Nguyen and McCurdy, 2015) |
| ggPm.At.leafepidermis.topview | *Arabidopsis thaliana* | Leaf | (Guo *et al.*, 2021) |
| ggPm.At.leaf.crossection | *Arabidopsis thaliana* | Leaf | (Tsukaya, 2013) |
| ggPm.At.seed.devseries | *Arabidopsis thaliana* | Seed | (Belmonte *et al.*, 2013) |
| ggPm.At.earlyembryogenesis.devseries | *Arabidopsis thaliana* | Embryo | (Wendrich and Weijers, 2013) |
| ggPm.At.shootapex.longitudinal | *Arabidopsis thaliana* | Shoot Apex | (Fuchs and Lohmann, 2020) |
| ggPm.At.inflorescencestem.crossection | *Arabidopsis thaliana* | Stem | (Shi *et al.*, 2021) |
| ggPm.Sl.root.crossection | *Solanum lycopersicum* | Root | (Ron *et al.*, 2013) |
| ggPm.At.leaf.topview | *Arabidopsis thaliana* | Leaf | (Vanhaeren *et al.*, 2015) |
| ggPm.At.rootelong.longitudinal | *Arabidopsis thaliana* | Root | (Shahan *et al.*, 2022) |
| ggPm.At.rootmatur.crossection | *Arabidopsis thaliana* | Root | (Shahan *et al.*, 2022) |
| ggPm.At.flower.diagram | *Arabidopsis thaliana* | Flower | (Taiz *et al.*, 2015) |
| ggPm.At.lateralroot.devseries | *Arabidopsis thaliana* | Lateral Root | (Torres-Martinez *et al.*, 2019) |
| ggPm.Ms.root.crossection | *Medicago sativa* | Root | Unpublished |

**Figure 4.**
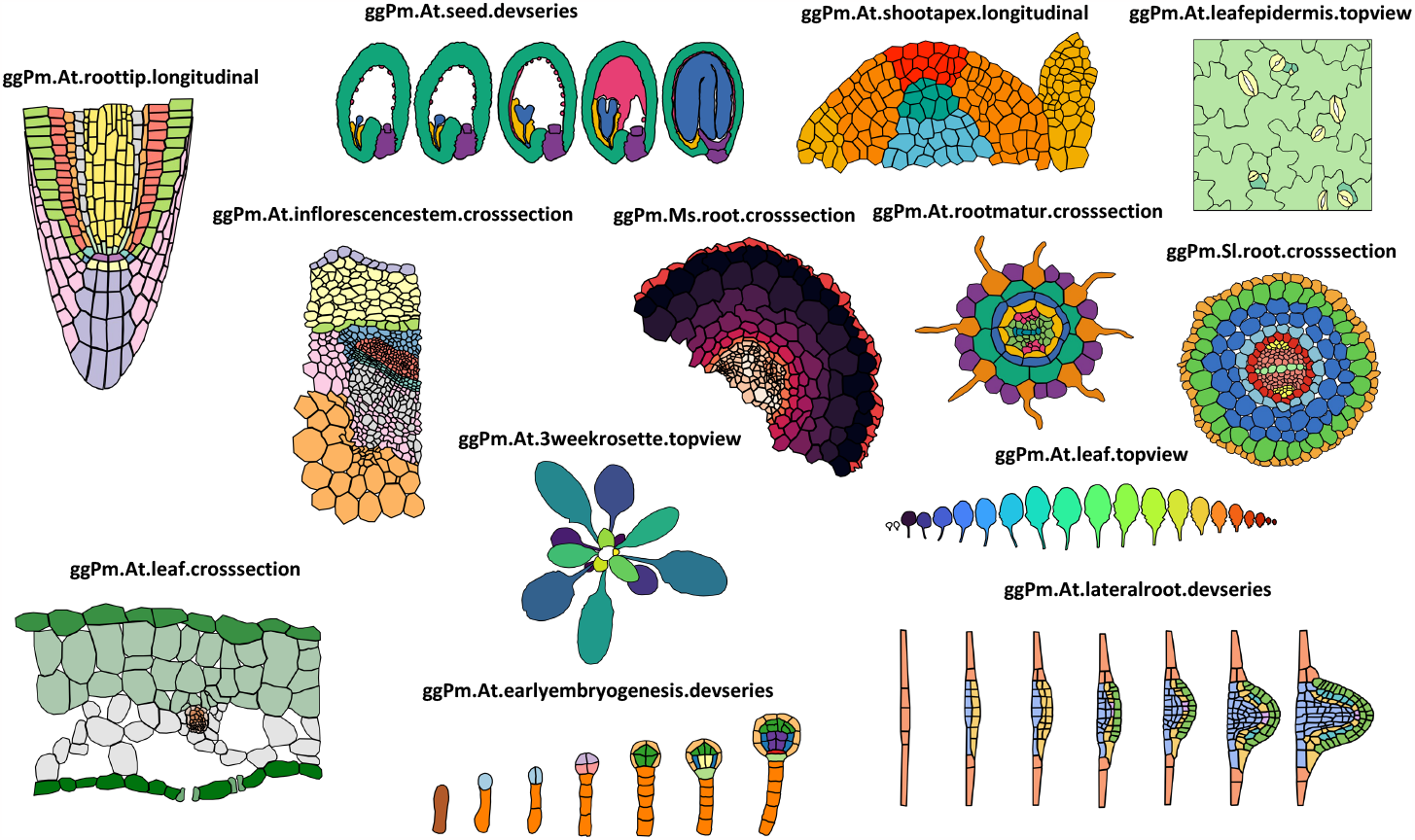
Examples of pre-loaded maps in the ggPlantmap package and their respective names. Pre-loaded maps were generated by tracing previously published plant images. Details of the maps, including the references, are given in Table 1 and can be retrieved from the ggPm.summary object in the package.

ggPlantmaps can be displayed graphically using the ggPlantmap.plot(), which is based on the well-known data visualization R package ggplot2 (Wickham, 2011). External quantitative data can be combined with table maps using the ggPlantmap.merge() function and displayed as heatmap with the ggPlantmap.heatmap() function.

### Tutorial for ggPlantmap

Installation instructions of ggPlantmap are provided in the <u>package github page</u>. We also included an extensive and detailed <u>user guide</u> through all the available functionalities of ggPlantmap. Additionally, we created a <u>walkthrough document</u> to guide users on creating their own ggPlantmap. We strongly recommend users to begin with the tutorials available in the GitHub page. In the sections below we will provide a tutorial to navigate through the package after it has been loaded to R (as described on the GitHub page). The files used for the tutorial can be found in the supplemental dataset 1.

#### Creating a new ggPlantmap

Creating a new ggPlantmap is fairly simple (Figure 5). First, open the provided plant image (“sample.jpg”, Supplemental dataset 1) in the open-source software Icy (https://icy.bioimageanalysis.org/). The creation of ggPlantmaps does not require high resolution images and can be performed with most standard image file formats (.jpg, .jpeg, .png, .tif, and others) as well as high-resolution microscopy files (.czi, .lif, and others). Second, trace individual compartments as individual Region of Interests (ROIs) using the polygon tool (Figure 5A). Upon tracing, it is important to name individual ROIs with specific names that will be used to distinguish the different levels (here: cell types) for color mapping. In the example provided in Figure 5A, we named cells in the most outward cortex layer of a *Medicago sativa* root cross section as ‘C1’ and cells in the second cortex layer as ‘C2’ (Figure 5A). Upon finalizing the tracing and naming of individual ROIs, select them all and save them as an XML file (Figure 5B). For this tutorial, we provided a finalized XML file (sample.xml) from the tracing of the provided sample image (Supplemental dataset 1). In R, this XML file can be converted into a ggPlantmap table object by using the XML.to.ggPlantmap() function as shown in Figure 5C. As mentioned above, the table consists of a column that describes the levels of ROIs (ROI.name), a column that contains a unique number identifier of individual ROIs (ROI.id), a column with the numeric order of individual points of the ROI shape (point), and the x and y coordinates of each point in a cartesian space (x,y).

**Figure 5.**
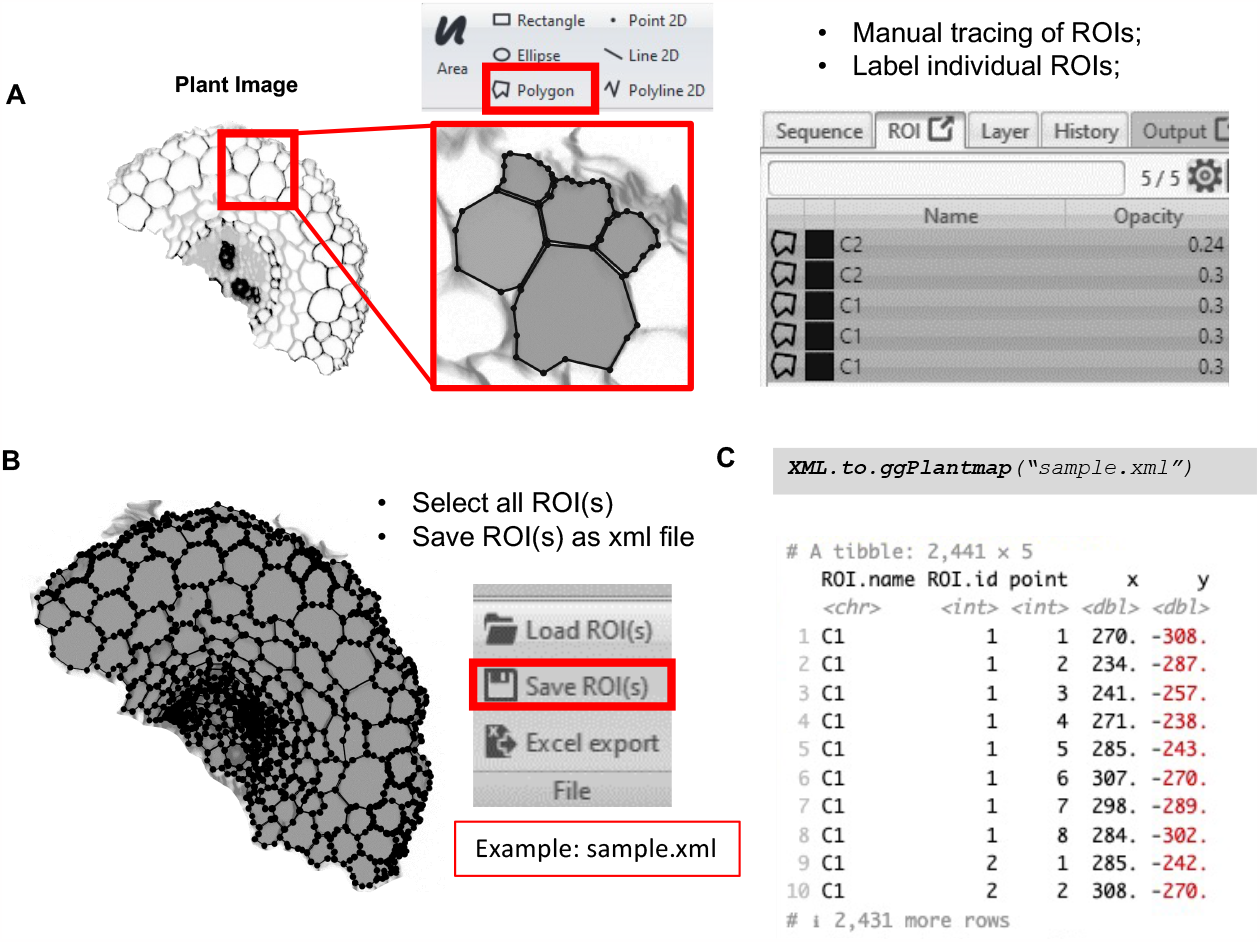
Creating your own map table. **(A)** In the Icy software, trace the individual structures using the ROI polygon tool. Individual ROIs need to be named with specific names that will be used for color mapping. Here, we used C1 for the cells in the outermost cortex layer and C2 for the cells in the second cortex layer. **(B)** After finalizing the manual tracing, save ROIs into an XML file. The time spent tracing images is dependent on the user experience and amount of detail in the image. The tracing of the example image here was performed in approximately one hour. **(C)** Use the XML.to.ggPlantmap() function to convert the stored ROIs into a ggPlantmap table object.

#### Plotting a ggPlantmap

We built the package with the flexibility of the well-known data visualization package ggplot2 (Wickham, 2011). With the ggPlantmap.plot() function, users can feed their newly created map objects or pre-loaded map tables into a ggplot2 plot object (Figure 6A). Built-in ggPlantmap.plot() function allows to further customize the plot. For example, users can select if they want to show the legends using the argument show.legend=FALSE (TRUE is the default) (Figure 6B) or change the line thickness of the plot by adjusting the linewidth argument (linewidth=1 is the default) (Figure 6C). Users can provide the column name to be used for color mapping (layer=ROI.name is the default) (Figure 6D). In the example provided in Figures 6D-E, color mapping was adjusted to respectively depict the zone and layer organization of the Arabidopsis shoot apical meristem pre-loaded map (ggPm.At.shootapex.longitudinal). Because ggPlantmap.plot() function is built using ggplot2, new lines can be added to further customize the display of ggPlantmaps using the ggplot2 logic (Figure 6F).

**Figure 6.**
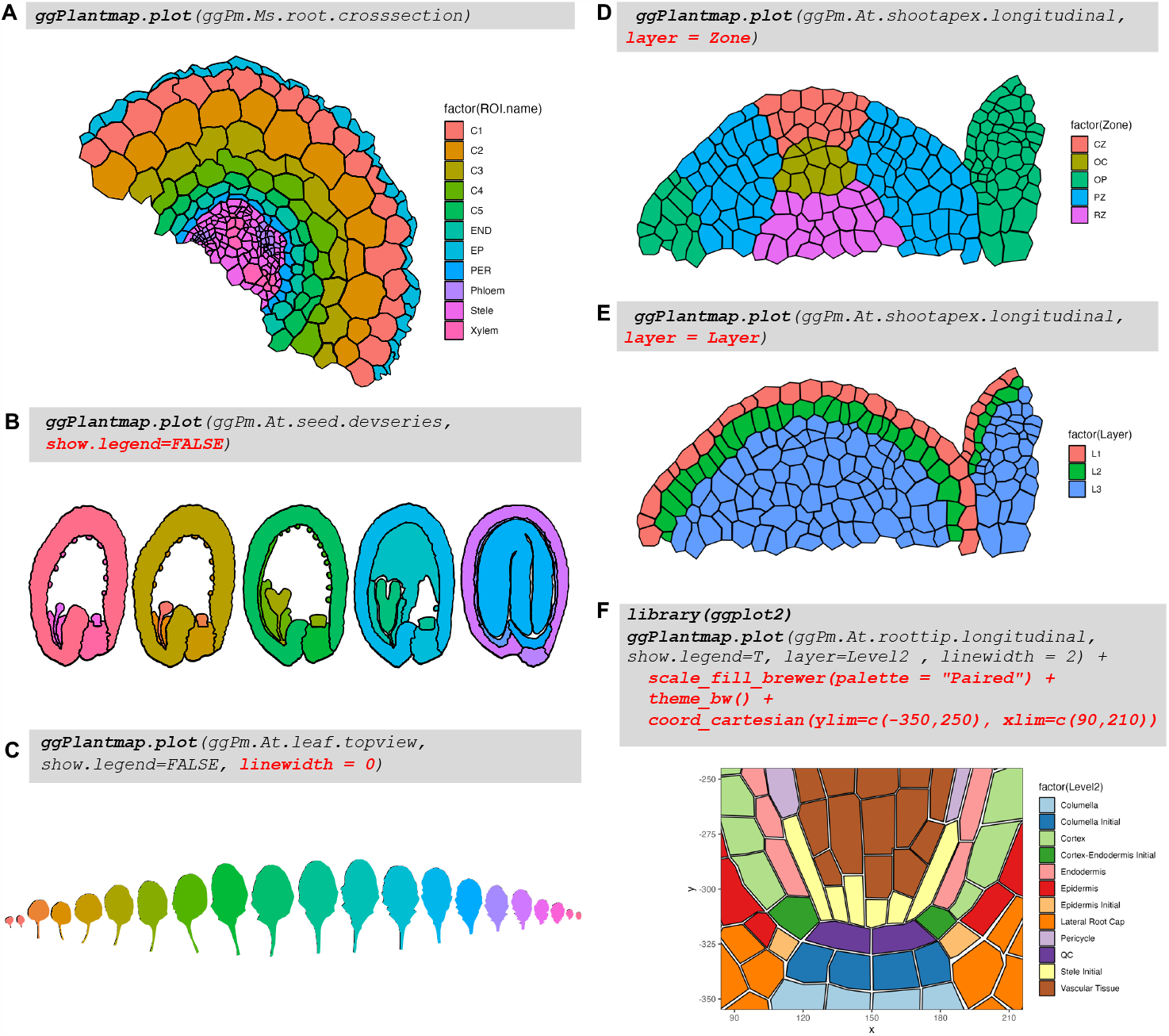
Example plot outputs from the ggPlantmap.plot() function. **(A)** Plot of the Medicago sativa root cross section pre-loaded map with default plotting options. **(B)** Removing the legends of plot of the Arabidopsis seed development series pre-loaded map with the argument show.legend=F. **(C)** Removing the outline of the Arabidopsis leaf pre-loaded maps with the argument linewidth=0. **(D&E)** Changing the color mapping argument (“layer”) to Zone (D) or Layer (E) column for the Arabidopsis shoot-apical meristem pre-loaded map. **(F)** ggPlantmap is build using ggplot2. Therefore, ggPlantmap.plot() function can be combined with multiple ggplot2 customizable options.

#### Integrating external quantitative data

The main goal of the ggPlantmap package is to give the user the ability to easily generate representative maps of large-scale quantitative datasets. We pre-loaded a sample quantitative data in the package (*ggPm*.*At*.*seed*.*expression*.*sample*). First, assign the expression data into a new object (expression.data <-ggPm.At.seed.expression.sample) and combine it with the ggPm.At.seed.devseries pre-loaded map using the ggPlantmap.merge() function (Figure 7). Merging both datasets requires specifying the key column names that contain the levels (ROI names) that match between datasets (id.x = “ROI.name” is the default). If key column names between datasets are not the same, indicate both column names in the id.x and id.y arguments of the function (id.x = id.y is the default). The output of this function is a new table with the quantitative values assigned for each one of the unique levels of the ggPlantmap. This new table can be used in the ggPlantmap.heatmap() function to generate a quantitative representative map as a heatmap (Figure 7). For producing the heatmap, the column name containing the quantitative values needs to be specified in the value.quant argument (Figure 7). Just like the ggPlantmap.plot() function, the ggPlantmap.heatmap() function can also be customized with ggplot2 arguments (Figure 7).

**Figure 7.**
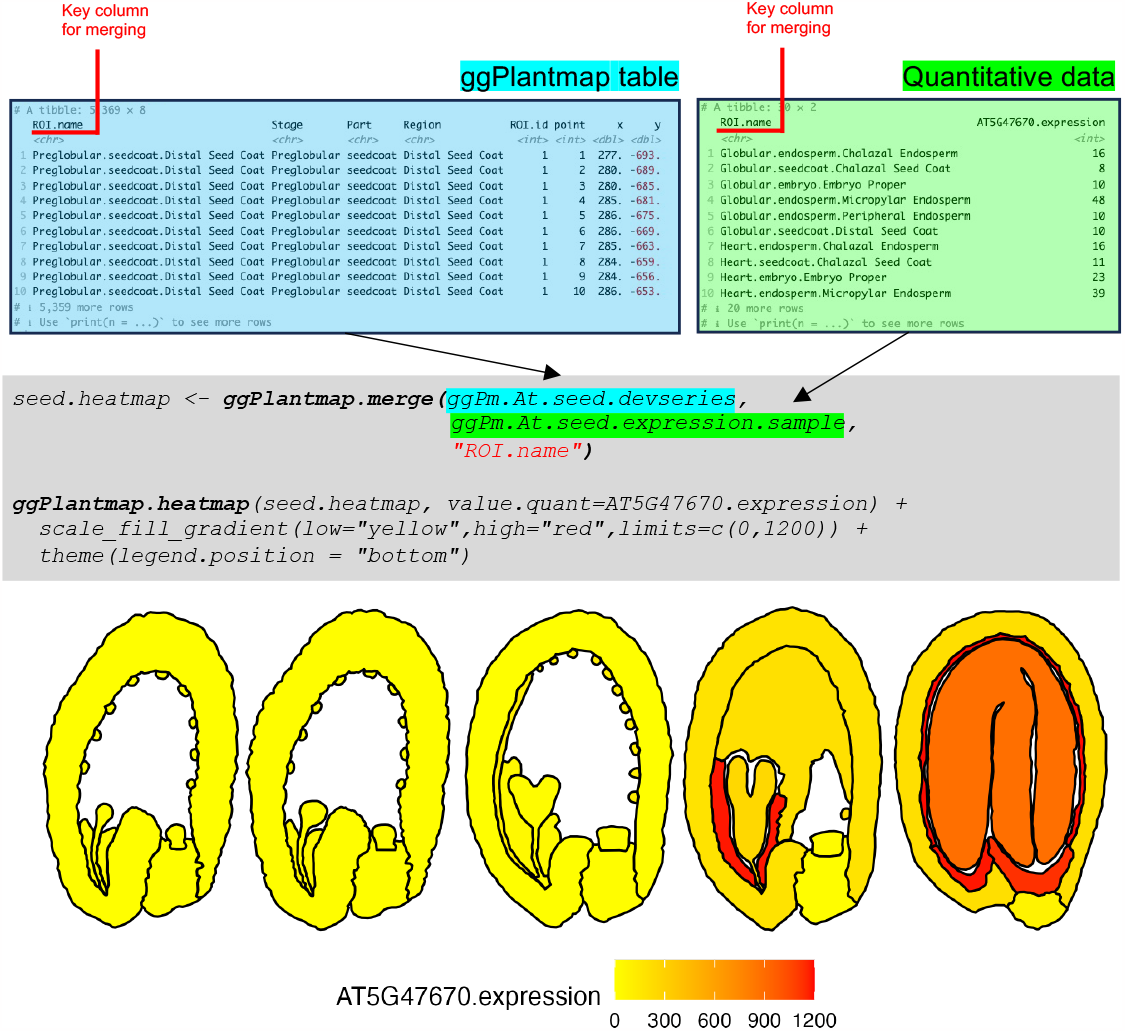
Integration of quantitative data into ggPlantmap. External quantitative data (green) can be combined with ggPlantmap tables (blue) using the ggPlantmap.merge() function. In the example provided, the expression of the *LEAFY-COTYLEDON1* (*LEC1, AT5G47670*) is integrated into the Arabidopsis seed development series pre-loaded map (ggPm.At.seed.devseries). The pre-loaded expression sample data (ggPm.At.seed.expression.sample) depicts the microarray signal of the *LEC1* gene in the microarray dataset of laser-capture microdissected tissues of the Arabidopsis seed (Belmonte *et al*., 2013). For simplicity, sample names in the dataset have been modified to match the ones found in the pre-loaded map. The third argument (in red) of the ggPlantmap.merge() function is the key column name that contains the matching ROI names between datasets. The output of the ggPlantmap.merge() function can be loaded into the ggPlantmap.heatmap() function to produce a heatmap that shows the quantitative expression of *LEC1* in the different compartments during the Arabidopsis seed development. As shown, this plot can also be customized with different ggplot2 options.

The generation of ggPlantmap heatmaps can be a powerful tool for the visualization of cell-type-specific quantitative profiles in large plant datasets. As an example, we implemented and generated quantitative representative maps using two publicly available cell-type-specific datasets: the microarray dataset for laser-capture microdissected tissues of the Arabidopsis seed (Belmonte *et al*., 2013) and the single-cell RNA-seq dataset for Arabidopsis root (Denyer *et al*., 2019) (Figure 8). With ggPlantmap, users can now streamline and generate maps to evaluate the expression of maps for their own specific datasets. In addition, ggPlantmap can be included in R/shiny interactive dataviewers for the better communication of large and complex datasets.

**Figure 8.**
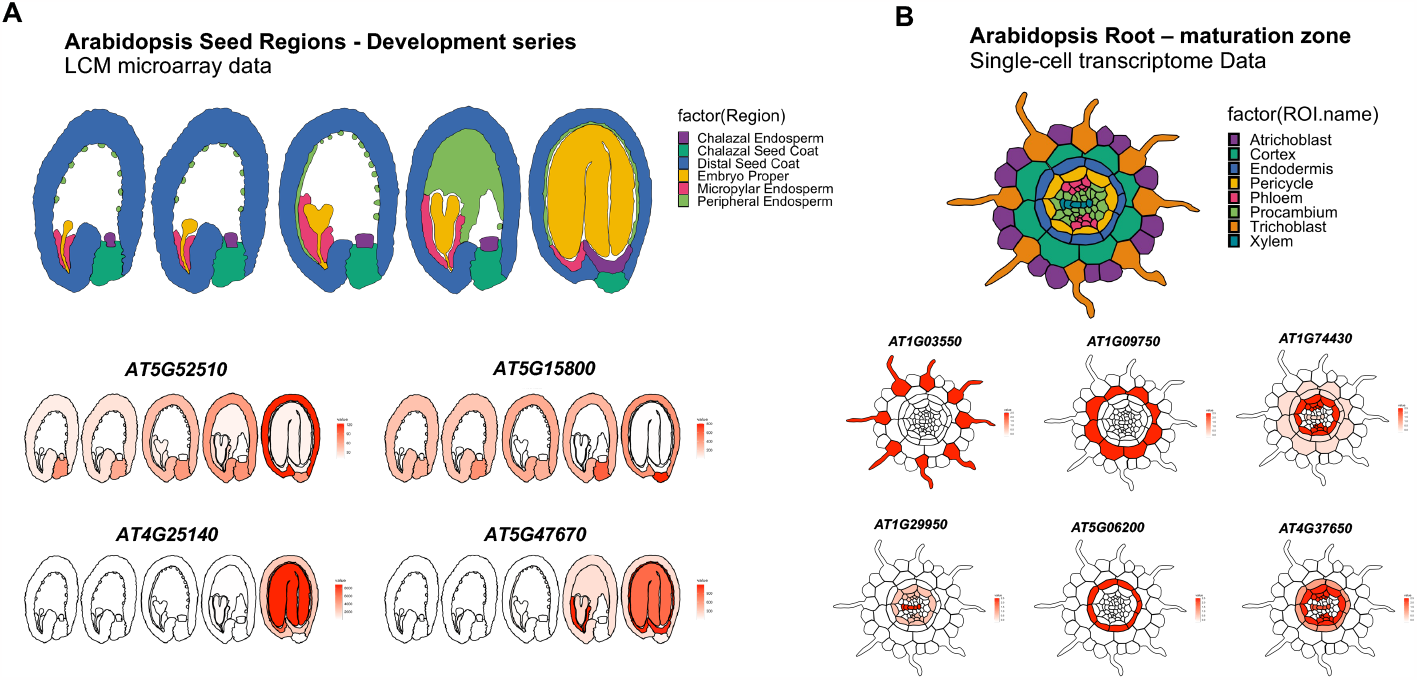
Using ggPlantmap to show tissue/cell-type-specific gene expression. We used ggPlantmap to show the tissue/cell-type-specific gene expression using two publicly available datasets. **(A)** Microarray dataset for laser-capture microdissected (LCM) tissues of the Arabidopsis seed (Belmonte et al., 2013) and **(B)** the single-cell RNA-seq dataset for Arabidopsis root (Denyer et al., 2019). Names of tissues and/or cell types in each specific dataset was changed to match the ones found in the ggPlantmap pre-loaded maps.

## Conclusion and Perspectives

Similar to a Plant eFP-browser, we built this package with a goal of displaying spatial profile of gene expression in unique plant tissues. However, we believe that the accessibility and simplicity of the package opens exciting possibilities for creative applications in plant biology beyond gene expression analysis (Figure 9). Its capacity to convert plant images into graphic maps that can be customized and worked in an R environment offers a flexible platform that can be used to communicate different types of quantitative data on plant images. We provide some conceptual examples in Figure 9. We envision ggPlantmap enabling intuitive plotting of quantitative data describing various cell or organ measurements (e.g. area, thickness, packing), concentrations (e.g. metabolites, nutrients, hormones), physiological measurements (e.g. conductivity, water potential), fluorescent reporters (e.g. calcium, auxin, oxygen), or fluorescent dyes (e.g. cell wall components). We encourage users to explore the full potential of this tool, adapting it to their unique research questions and leveraging its capabilities to present and interpret a broad spectrum of biological quantitative data.

**Figure 9.**
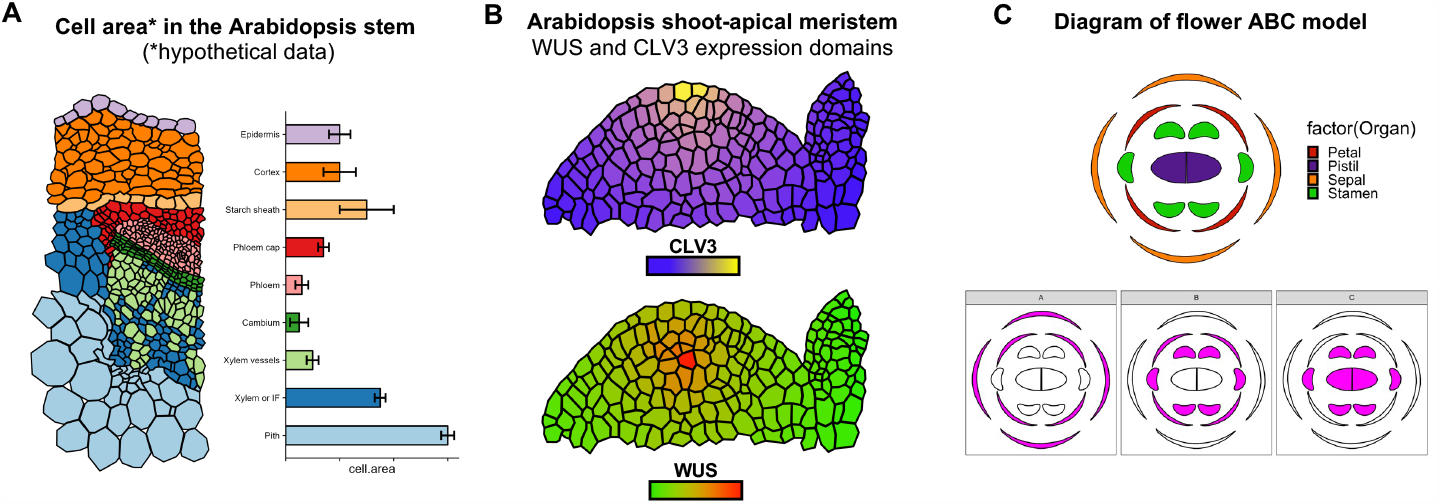
Examples of potential application of ggPlantmap for the communication and education of plant sciences. **(A)** Combining ggPlantmap with different plots for the better communication of quantitative data in plant biology. In the example provided, we show hypothetical data of cell area of different cell types of the Arabidopsis stem by pairing ggPlantmap with a bar chart. **(B)** Using ggPlantmap to show the *CLAVATA3* (*CLV3*) and *WUSCHEL* (*WUS*) expression domains in the Arabidopsis shoot-apical meristem. **(C)** Using ggPlantmap to show the ABC model of flower development in Arabidopsis.

Additionally, we believe that this package offers a valuable tool for education and communication in plant sciences to a broader audience. By converting complex quantitative data into visually accessible formats, it facilitates the communication through engaging visuals for educational settings and enhances communication with the public. Its user-friendly features make it a powerful asset for educators and science communicators, bridging the gap between the scientific community and a broader audience.

The ggPlantmap package is an open-source project and we encourage the plant research community for contributions and creation of maps that will be continuously loaded into the package. In the package github page, we included <u>instructions</u> on how newly community generated maps can be included in the package. We hope that ggPlantmap can play an important role in the data visualization toolbox by offering an open, accessible, and customizable solution for creating quantitative image maps from plant images. By providing plant researchers with the means to independently generate maps from plant images, this package will empower plant scientists to own the means of exploring their data in creative and exciting ways. We encourage the plant research community for feedback and suggestions through the corresponding emails or via the github page.

## Acknowledgements

We thank Andres Romanowski, Kyra van der Velde, Lisa Oskam, Monica Garcia Gomez and Pierre Gautrat for suggestions and trials of the initial versions of the package. We also acknowledge Dorota Kawa and Kirsten H Ten Tusscher for reviewing this manuscript.

